# *Ex vivo* modelling of lung tissue resident antimicrobial responses

**DOI:** 10.1101/2025.01.27.634989

**Authors:** Hélèna Choltus, Julien Prados, Niccolo Bianchi, Nelli Heikkila, Wolfram Karenovics, Veronique Serre Beinier, Nina Leitner, Benoit Bédat, Fedor Bezrukov, Beryl Mazel-Sanchez, Virginie Prendki, Alexander Lobrinus, Christiane Eberhardt, Simone Becattini, Mirco Schmolke

## Abstract

Tissue resident host responses to microbial infections in the respiratory tract are highly dynamic in space and time and rely on the interaction of a multitude of cell types. In an attempt to model these multicellular responses reliably in cell culture, we compare here the global transcriptional antimicrobial response to infection with influenza A virus (IAV) in precision cult lung slices (PCLS), volume defined organ discs largely maintaining the cellular composition and 3D architecture of the donor lung. To permit a fair comparison of host responses in an isogenic background we first challenged mice in vivo and murine PCLS (mPCLS) and assess host transciptomic changes by unbiased RNAseq. While core antiviral responses overlapped substantially, mPCLS lacked certain features—such as type II interferon expression—likely due to the absence of infiltrating immune cells responses. Importantly, when expanding our findings to immune experienced human precision cut lung slices (hPCLS), we find a much broader antiviral response after IAV challenge, including type I, II and III interferons, suggesting the presence of responsive tissue resident lymphocytes. To prove specificity of this response we infected hPCLS with Streptococcus pneumoniae. Ex vivo tissues responded with a distinct proinflammatory gene profile including IL1A, IL1B and IL17 expression. Blocking of IL-1 signaling partially inhibited the proinflammatory response, suggesting cellular cross-talk and a complex and specific antimicrobial reaction in this *ex vivo* model. In conclusion diversified tissue resident immune cell compartment distinguishes the human *ex vivo* model, making it an ideal system for microbiological and immunological research.

**Importance:** Pathogen interactions with the lung are very dynamic processes. In biomedical research it is paramount to model these processes in the laboratory as accurately as possible. Influenza A virus has been extensively studied in epithelial cell culture models, including advanced organoids and organ on a chip systems. We use here ex vivo cultured PCLS and use transcriptomics to assess the global tissue resident host response to viral and bacterial challenge. Our data show 1) that murine PCLS faithfully reflect core responses to viral infection, while missing proinflammatory responses linked to infiltrating immune cells and 2) that human PCLS show a highly diversified tissue resident immune response to viral infection due to previous exposures of the host to this pathogen. These responses are clearly distinct from antibacterial gene profiles. Our data advertise PCLS as a complex and realistic model to study tissue resident immune responses to microbes in a human system.

## Introduction

Respiratory infections with viral or bacterial pathogens are the 5^th^ leading cause of death worldwide and the leading cause of death in low income countries (excluding COVID-19) according to 2021 WHO estimations^1^. Continuous exposure to the surrounding air makes the lung mucosa an ideal entry port for airborne viral and bacterial pathogens. Influenza viruses alone are responsible for more than half a million deaths annually often complicated by secondary bacterial pneumonia, mostly with Streptococcus pneumoniae or Staphylococcus aureus. A major initial barrier against establishment of respiratory infections is the antimicrobial host response elicited by respiratory epithelial cells in conjunction with tissue resident immune cells. Modelling the complexity of this first line of defense in cell culture setting remains however challenging, in part since the human lung consists of more than fifty cell types^2^. 2D cell culture models do not reflect this cellular diversity and the complex spatial 3D organization of respiratory tissue.

Precision cut lung slices (PCLS) are *ex vivo* cultured, i.e. volume defined discs of human or animal lung tissue origin and bridge conventional cell culture models and *in vivo* systems ^3,4^. They preserve the cellular composition and 3D architecture of lung tissue and are arguably the most complex *ex vivo* model to study lung physiology and pathology^5^. Since they are disconnected from the circulation of the host organism, they constitute an ideal scenario to study the tissue resident immune response to invading microbes. We compare here the tissue resident host response to viral and bacterial challenge in a murine and human ex vivo model with data from in vivo infected lung tissues.

## Results

The antiviral host response to influenza A viruses in the respiratory tract is an extraordinarily dynamic process initiated by infected epithelial cells and surveilling tissue resident immune cells. This first line of defense can be decisive for the outcome of infection. It establishes an antiviral state in the respiratory tissue, most importantly induced by cytokines of the interferon family. It further attracts immune cells from the blood and the interstitial space into the lung to limit and clear viral infection.

To delineate early tissue resident antiviral responses from those after immune cell infiltration, we turned to the mPCLS model (**Fig 1A**). Brightfield microscopy confirmed the maintenance of typical lung tissue architecture after tissue slicing (**Fig 1B**). mPCLS were cultured for several days without signs of tissue disintegration (**Fig. S1**). The immune cell compartment of mPCLS was predominantly characterized by naïve CD4 T cells and B cells in the lymphoid panel (**Fig. 1C, gating strategy in Fig. S2A**) and monocytes (classical and non-classical) in the myeloid panel (**Fig. 1D, gating strategy in Fig. S2B**). As compared to published in vivo data^6^, mPCLS show a slightly higher proportion of B lymphocytes *vs.* T lymphocyte, and a generally lower representation of myeloid cells, suggesting a that the latter might be partially excluded in the process of PCLS preparation.

**Figure 1:**
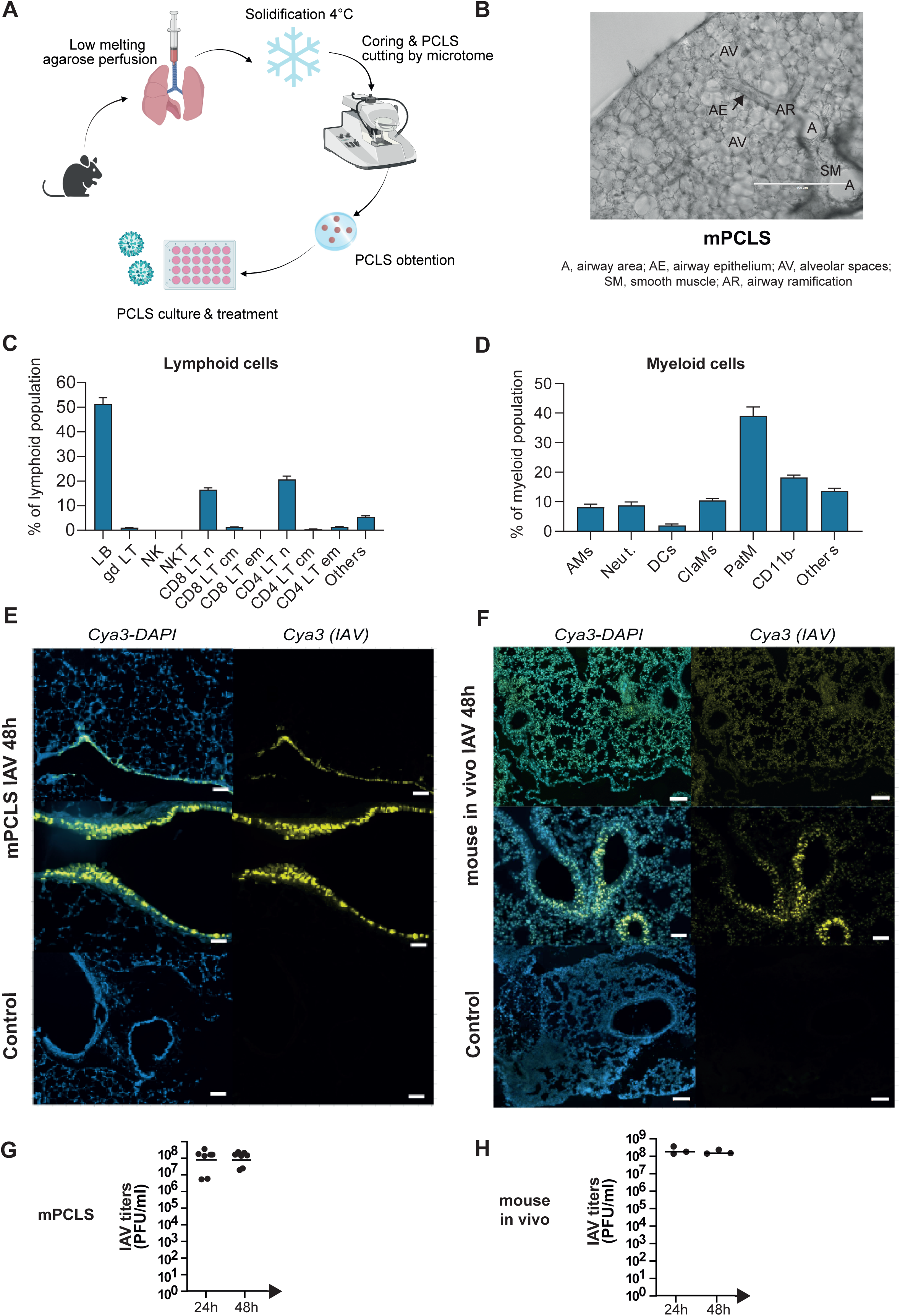
Modeling the early host response against Influenza A virus in murine PCLS. A) Schematic of PCLS generation: lungs are perfused with a low melting agarose solution to preserve the 3D lung architecture. After solidification at 4°C, lungs are cored into regular cylinders and processed to slicing using a microtome. PCLS are then kept in culture medium at 37°C. B) Brightfield picture of an untreated murine PCLS. Scale bar represents 400µm. C-D) Untreated PCLS were enzymatically digested one day after slicing to characterize baseline composition of lymphoid (C) and myeloid (D) immune cells. Results are presented in % of total lymphoid population and in % of total myeloid population (N=3). **Lymphoid cells**: LB: lymphocytes B, gdLT: γδ T lymphocytes, NK: natural killer cells, NKT: natural killer T cells, LT: lymphocyte T, n: naïve, cm: central memory, em: effector memory. **Myeloid cells**: AMs: Alveolar macrophages, Neut: neutrophils, DCs: dendritic cells, ClaMs: classical monocytes, PatMs: patrolling monocytes. Mice and mPCLS were respectively infected with 10^4^/10^5^ PFU of IAV (A/Netherlands/602/2009/H1N1) for 24 or 48h. E-F) In situ RNA hybridization of viral nucleoprotein (Cyanine 3) was used to visualize IAV at 48hpi in mPCLS sections (E) and in vivo in mouse lungs (F). Confocal observation was done at x100 (top pictures IAV & uninfected, size bar represents 100µM) or at x200 (bottom pictures, size bar represents 50µm). G-H) Titers were measured by plaque-assay on mPCLS (G, N=4, duplicates) and on in vivo infected lung (H, N=3) at 24 and 48hpi.

The porous nature of this *ex vivo* model permitted IAV infection with a clinically relevant H1N1 strain influenza A/Netherlands/602/2009 (H1N1) throughout the tissue as demonstrated by specific RNAscope staining of vRNA in mid-sections 48h post infection (**Fig. 1E**). We did however observe a preference for infection in larger airways. A similar distribution of vRNA signal was found in *in vivo* infected lung tissue 48h post infection (**Fig. 1F**). In line with this viral titers *ex vivo* and *in vivo* reached comparable levels 48h post infection (**Fig 1G and H**).

Using a bulk RNAseq approach we compared transcriptional responses in IAV infected lung tissue *in vivo* and *ex vivo*. Not surprisingly the PCA separated host responses clearly between the two models (along PC1), while the IAV induced parallel shift of the global transcriptome along PC2 in the two systems, which was less prominent especially in the *ex vivo* model (**Fig. 2A**). Accordingly, we found substantially more differentially regulated genes (DEG) *in vivo* than in mPCLS (**Fig. 2B-D**). Approximately 20% of the upregulated protein coding DEG were classified as “immune system” related in the Reactome database^7^ (**Fig. 2B**). Notably, IAV induced DEG in mPCLS were largely found upregulated *in vivo* (**Fig. 2B**). The two gene sets correlated with a R^2^: 0.5 (**Fig. 2E**). An analysis of transcription factor binding sites in promoters of upregulated genes in mPCLS indicated that IRF3 and STAT2 are mostly responsible to drive the *ex vivo* response, while *in vivo* NFκB, STAT5B, NOD2 and IRF3 were the most significantly enriched transcription factors (**Fig. S3A and S3B**)^8–10^. Comparative Reactome analysis of mPCLS and *in vivo* mouse responses to IAV revealed a strong enrichment in immune system including cytokine signaling, innate and adaptive immunity in both models (**Fig. 2F**). While for mPCLS, the response was restricted to immune system activation, we found *in vivo* additionally an enrichment in hemostasis, extracellular matrix organization and programmed cell death among the upregulated DEG (**Fig. S3C-E**).

**Figure 2:**
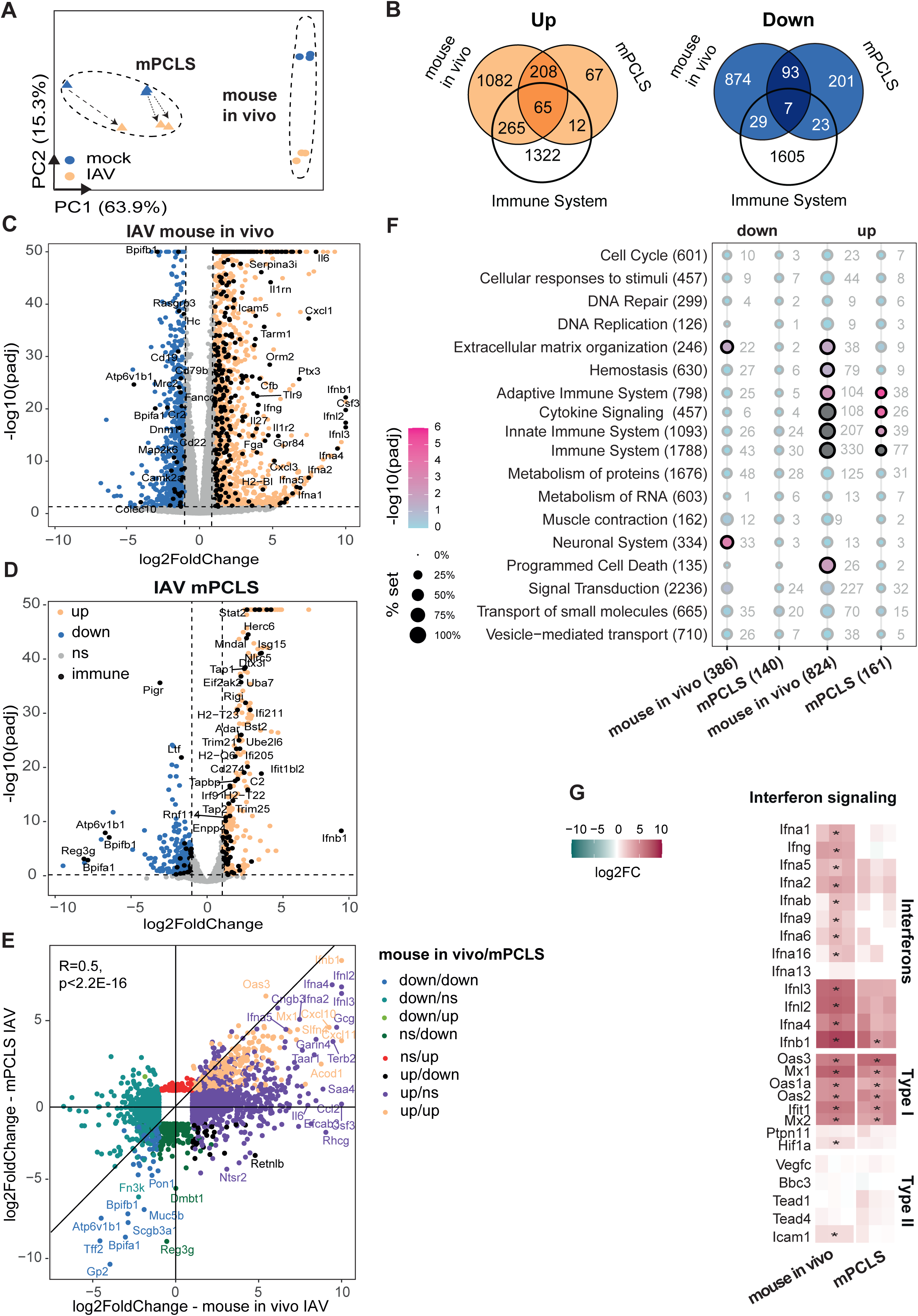
Comparative transcriptomic analysis of the early host response to Influenza A virus in murine lung in vivo and ex vivo. Mice and mPCLS were respectively infected with 10^4^/10^5^ PFU of IAV (A/Netherlands/602/2009/H1N1) for 48h. Tissue was processed to bulk RNA sequencing and host response was analyzed on R using Deseq2 package. A) Principal component analysis illustrates the transcriptional response of mock treated (blue) and IAV-infected lung tissue (orange) in vivo and in mPCLS. Arrows indicate the pairs of mock and infected PCLS from the three independent experiments. B) Venn diagrams of the commonly upregulated (orange) or downregulated (blue) genes in mouse in vivo and mPCLS infected with IAV. The white circle indicates the genes related to immune system pathways (using Reactome annotations). C-D) Volcano plot showing differentially expressed genes between IAV-infected and mock samples in mouse in vivo (C) and in mPCLS (D). Log2 fold-change induction is indicated in the x-axis, -log10(p-adjusted) in the y-axis. Genes have been represented in: orange (upregulated), blue (downregulated), grey (not significant). Genes associated to immune system pathways are depicted in black. E) Scatter plot comparing the genes regulated by IAV in mouse in vivo (log2 Fold Change in x-axis) and in mPCLS (log2 Fold Change in y-axis). Correlation is indicated with R Pearson coefficient (R=0.5, p<2.2E-16). Color legend indicates the direction for mouse in vivo/mPCLS genes regulated. F) Pathway enrichment analysis was conducted using hypergeometric tests against Reactome.org database. -log10 (p-adjusted) is given by a color scale. Coverage percentage of the pathway set is indicated by the size of the dot. Pathways are indicated with the total number of genes in brackets and the number of genes significantly regulated are indicated next to the dot. Significantly enriched pathways are indicated by a black circle. G) Heatmap illustrates the regulation of interferon signaling genes by IAV 48hpi in mouse in vivo and in mPCLS. Genes are represented by the log2 Fold Change (pink: positive log2FC, blue: negative log2FC), * indicates if the gene is significantly regulated by IAV infection.

A common denominator of the antiviral response in human and murine hosts is the interferon response^11^. *Ex vivo* type I and III interferon genes were generally upregulated to a much lower extend than *in vivo*, with IFNB1 being the most highly induced type I interferon gene (**Fig. 2G**). Accordingly, we found upregulation of typical ISGs like MX1, OAS1, or IFIT1 ((**Fig. 2G**) amongst others). Notably, mPCLS were completely devoid of a type II interferon response (**Fig. 2G**). Accordingly, ICAM1 an experimentally confirmed type II IFN dependent gene^12^ was not upregulated in mPCLS after IAV challenge but type I/type III dependent genes were. IFNγ is almost exclusively produced by lymphoid cells (NK, NKT, effector/memory CD4^+^, CD8^+^ and γδ T lymphocytes). These cells were largely absent in mPCLS (**Fig. 1C**) possibly explaining the lack of IFNγ in response to IAV infection.

Both our PCLS donor mice and the *in vivo* infected animals were immunologically naïve to IAV infection, explaining the lack of tissue resident memory cells. To simulate tissue responses of immune-experienced mice we generated mPCLS from IAV (A/Netherlands/602/2009 H1N1) preinfected mice one month after primoinfection (**Fig. 3A**) and challenged these immune experienced mPCLS with the same virus strain *ex vivo*. The presence of memory T lymphocytes was confirmed by FACS (**Fig 3B and gating strategy in Fig S4**). Viral replication was reduced in *ex vivo* tissues from IAV primed mice as confirmed by qPCR for vRNA and by titration of viral supernatants (**Fig. 3C and 3D**). Surprisingly, this did however not result in a generally heightened transcriptional response in IAV primed murine lung tissues *ex vivo* as assessed by targeted qRT-PCR (**Fig. 3E**). Our data hence suggest an increase in viral defense with improved control of viral replication one month after priming, which does not generally depend on an upregulation of innate immune genes, nor seems to promote it upon *ex vivo* reinfection of tissues.

**Figure 3:**
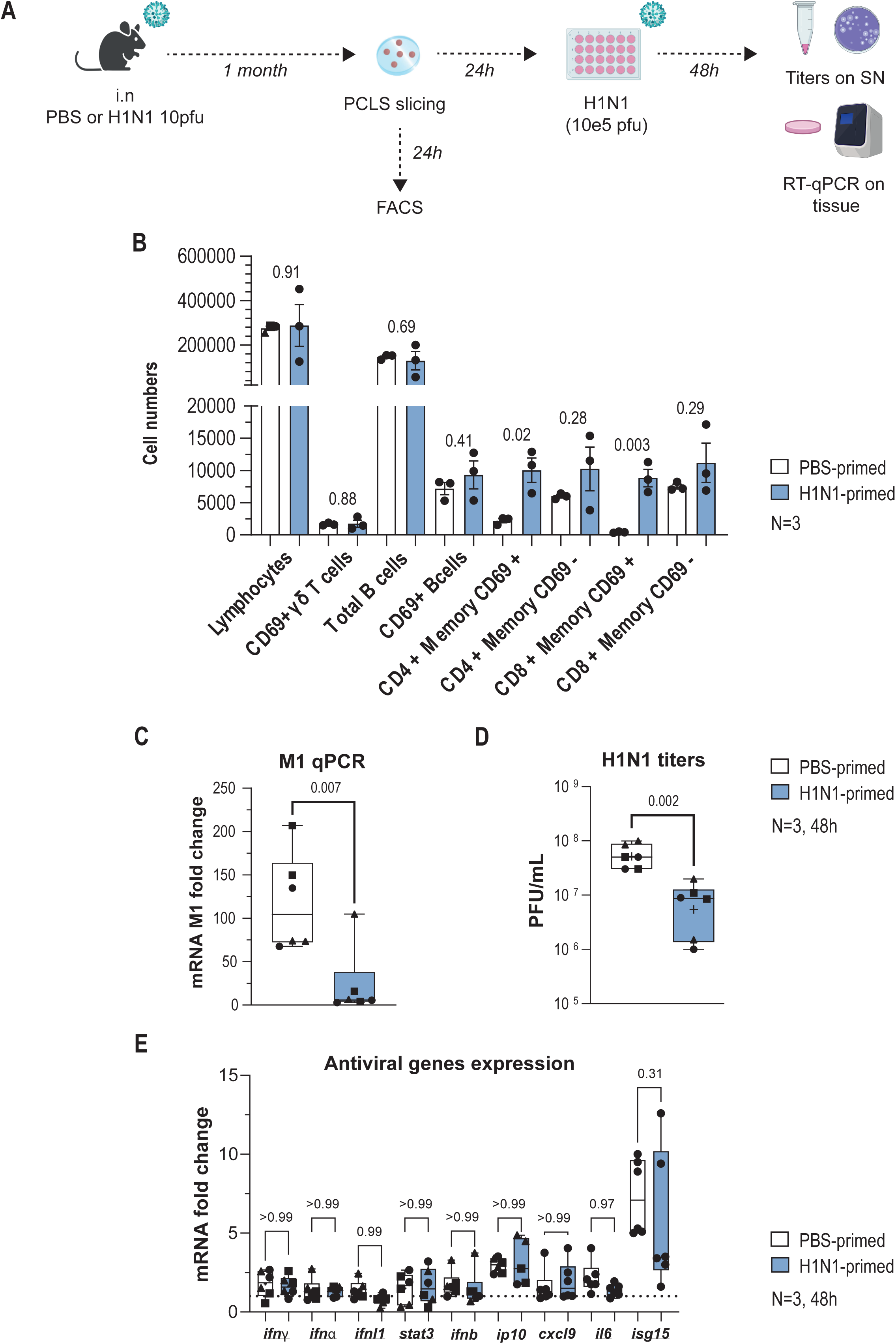
Characterization of the innate immune response in IAV-primed mPCLS. A) Mice were intranasally primed with PBS or 40pfu of A/Netherlands/602/2009/H1N1 (N=3, duplicates). One month later, mice were sacrificed and mPCLS were generated and homologously re-infected with 10^5^ pfu of IAV for 48h. Supernatants and tissues were collected for titers and RNA extraction. B) Flow cytometry was conducted on untreated mPCLS from PBS or IAV-primed mice. Lymphoid cell population was quantified. C) Viral replication was measured at 48hpi by RT-qPCR on M1 gene on IAV-infected PCLS from PBS and IAV-primed mice. D) Viral titers 48 hpi were measured by plaque-assay. E) RT-qPCR on antiviral genes was performed 48hpi. Fold change over mock are represented. Statistical analysis was done with a t-test (N=3, each symbol represents one independent experiment). PBS-primed mPCLS are represented in white boxes and IAV-primed mPCLS in blue boxes. p-value is indicated.

In average a human is exposed every 5-10 years to a new influenza virus^13^. These repeated infections lead to an accumulation of tissue resident memory T cells in adults. In elderly influenza specific _trm_T-cells decline with advanced age in lung tissue, which could in part contribute to the reduced antiviral response^14^. Since elderly form a major population at risk for severe IAV infection, we asked if PCLS from an immune experienced elderly human donor would provide a more realistic model of tissue resident immune responses against IAV.

PCLS were cut from histologically healthy bystander tissue of tumor resections (for patient cohort data **see Table S1**) and maintained typical 3D architecture of human lung tissue (**Fig. 4A**). Notably, airspaces were much larger than those found in mPCLS (**Fig. 1B**). *Ex vivo* hPCLS were cultured for several days without signs of cell death comparable to mPCLS (**Fig. S1**). In contrast to murine tissues T lymphocytes were the dominant lymphoid cell population in hPCLS among these more than 50% effector memory CD4 and CD8 T cells (**Fig. 4B and gating strategy in Fig. S5A**). Additionally, we found low percentages of NK and NKT cells, absent in *ex vivo* mouse tissues. In the myeloid panel, monocytic myeloid derived suppressor cells ^15^ and neutrophils were dominant (**Fig. 4C and gating strategy in Fig. S5B**).

**Figure 4:**
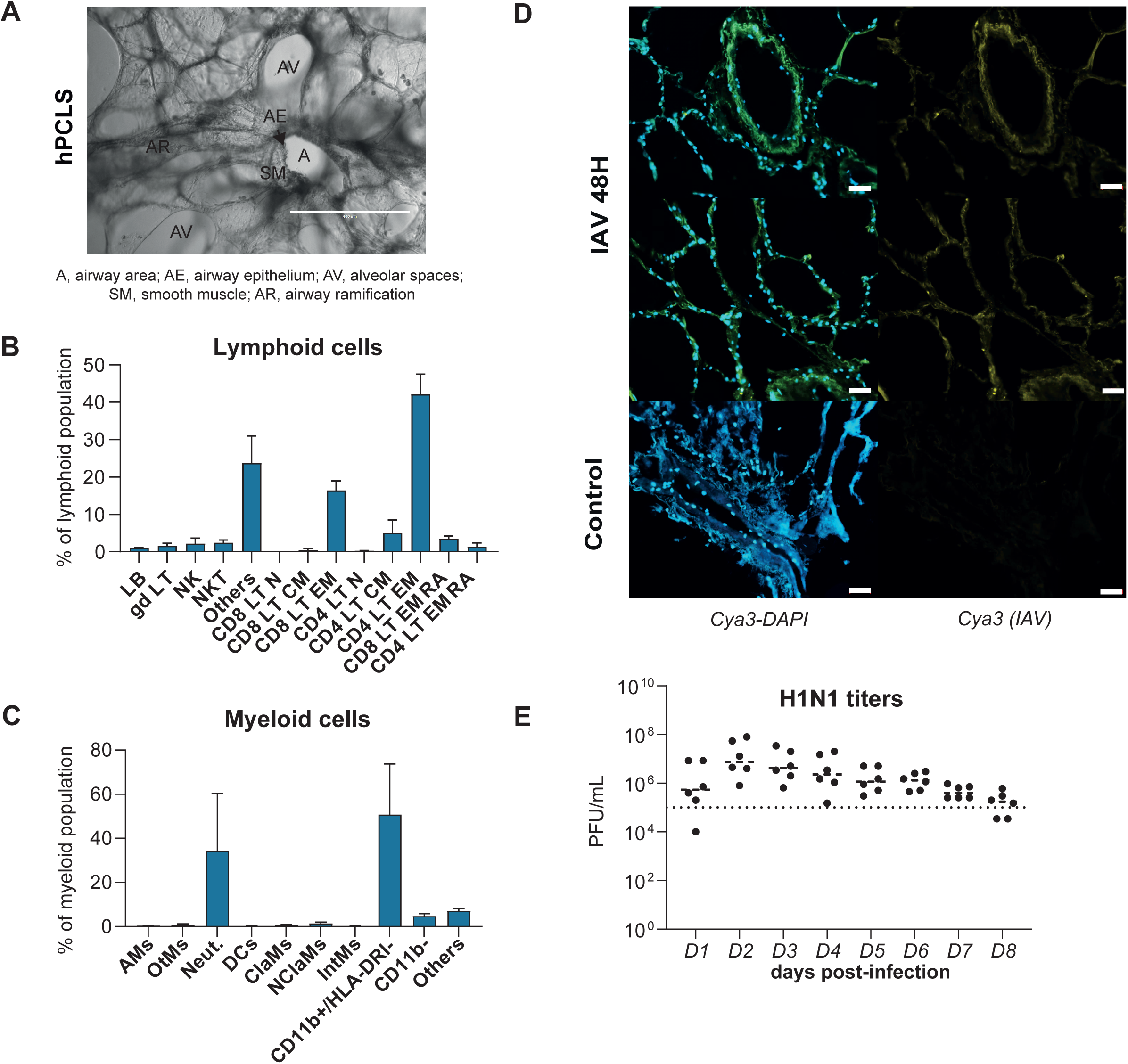
Antimicrobial host response to viral challenge in the human PCLS model. A) Brightfield picture of an untreated human PCLS in culture (magnification X100, size bar represents 400µM). B-C) Untreated PCLS from 3 patients were enzymatically digested one day after slicing to describe the baseline composition of lymphoid (B) and myeloid (C) immune cells. Results are presented in % of total lymphoid population and in % of total myeloid population. **Lymphoid cells:** LB: lymphocytes B, gdLT: γδ T lymphocytes, NK: natural killer cells, NKT: natural killer T cells, LT: lymphocyte T, n: naïve, cm: central memory, em: effector memory, emra: effector memory CD45RA^+^. **Myeloid cells**: AMs: Alveolar macrophages, OtMs: other macrophages, Neut: neutrophils, DCs: dendritic cells, ClaMs: classical monocytes, NClaMs: non-classical monocytes, IntMs: Intermediate monocytes. D) In situ RNA hybridization of viral nucleoprotein (Cyanine 3) was used to visualize IAV at 48hpi in hPCLS sections. Magnification x200. Size bar is representing 50 µm. E) hPCLS were infected with 10^5^ pfu of IAV (A/Netherlands/602/2009/H1N1) for 8 days, viral titers were measured daily by plaque-assay (3 patients in duplicates).

As for mPCLS, IAV penetrated the entire tissue (**Fig 4D**) and resulted in robust viral titers 48hpi (**Fig. 4E**). We found little donor to donor variation in the replication data, despite using tissues from different anatomical locations of the lung and diverse underlying disease states (**Table S1**), supporting the robustness of this model system for microbiological research. The PCA clearly separated mock treated from IAV infected hPCLS. This is similar to data obtained from an influenza virus RNA positive lung autopsie tissue from a patient who died shortly after an influenza B virus diagnosis when compared to freshly isolated healthy lung tissue material (**Fig. 5A**). In contrast to the mouse model, comparable numbers of DEG were upregulated in hPCLS and human *in vivo* infected lung tissue. However, at protein coding gene level we found only an small overlap of the upregulated DEG of hPCLS with the *in vivo* tissues (**Fig. 5B-C**). This could be the consequence of a number of confounding effects: comparison of non-isogenic hosts, comparison of an influenza A virus vs an influenza B virus infection, different sequencing protocols (standard illumina RNAseq *vs.* exon enrichment for *in vivo*) and most importantly, differences in the time point post infection. Nevertheless, the reactome analysis provided however substantial overlap in upregulated gene modules related to innate immune response, cytokine signaling and DNA replication, the latter being absent in murine samples (**Fig. 5D and E**). On gene level the intersection of upregulated genes contained known ISGs like (MX2, IFI6, IFIT1, IFIT5, IFIT3, IFI27, IRF27, IRF9, ISG15, IFITM1) were found. IAV infection robustly induced type I, II and III interferons in hPCLS. Upregulation of IFNγ and IFNγ-induced genes after IAV infection (**Fig. 5F**) is of interest in this model since it confirms the activation and functionality of tissue resident lymphoid cells.

**Figure 5:**
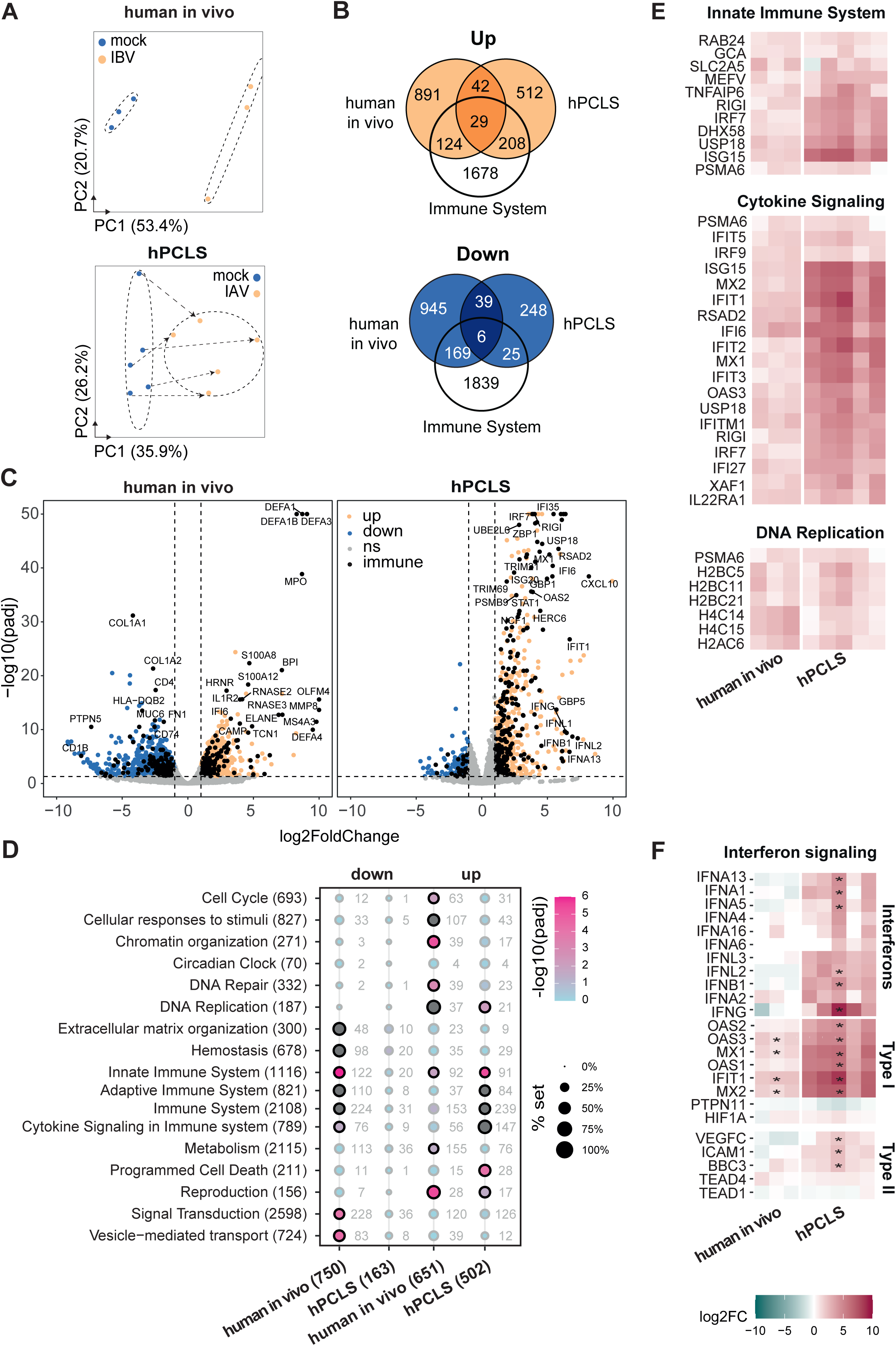
Antimicrobial host response to viral challenge in the human PCLS model *versus* in vivo human lungs. Human PCLS from five patients were infected with 10^5^ pfu of IAV (A/Netherlands/602/2009/H1N1) for 48h. In parallel, lung biopsies from a naturally Influenza-infected patient *in vivo* (triplicates from three different lobes) were compared to unprocessed lung samples from three uninfected patients. Bulk RNA sequencing was performed on these samples and analysis was conducted using Deseq2 package on R software. A) Principal component analysis illustrates the global transcriptomics response of control (blue) and Influenza-infected (orange) samples in human in vivo (left) and in hPCLS (right). B) Venn diagrams of the commonly upregulated (orange) or downregulated (blue) genes in human *in vivo* and hPCLS after influenza infection. The white circle indicates the genes related to immune system pathways. C) Volcano plot showing differentially expressed genes between Influenza-infected and mock samples in human in vivo (left) and in hPCLS (right). Log2 fold-change induction is indicated in the x-axis, -log10(p-adjusted) in the y-axis. Genes have been represented in: orange (upregulated), blue (downregulated), grey (not significant). Genes associated to immune system pathways are illustrated in black. D) Pathway enrichment analysis was conducted using Reactome database. -log10 (p-adjusted) is given by a color scale. Coverage percentage of the pathway set is indicated by the size of the dot. Pathways are indicated with the total number of genes in brackets and the number of genes significantly regulated are indicated next to the dot. Pathways significantly enriched are surrounded by a black circle. E) Heatmaps indicate the commonly up genes in human in vivo and in hPCLS after influenza infection for Innate Immune System, Cytokine Signaling and DNA replication pathways. Genes are ranked by the log2 Fold Change (upregulated: pink, downregulated: blue). F) Heatmap of genes associated with interferon signaling. The significance of regulation by IAV in human in vivo and in hPCLS is indicated by a *.

Next we were interested in testing the specificity of the transcriptomic response of hPCLS to a viral infection. Using a bacterial challenge with *S. pneumoniae* (serotype 3) we expected a distinct host response. As for the viral infection, bacteria penetrated the tissue to the center section **(Fig. 6A**). 24h post infection with *S. pneumoniae* we observed a substantial increase in bacterial titers associated with the lung tissue (**Fig. 6B**). The host response to Spn infection was showed a higher donor variation and was generally less pronounced as for the viral challenge (**Fig. 6C**), with patient 1 not responding to the bacterial challenge despite robust bacterial replication. Since the same patient tissue reacted well to viral challenge (**Fig. 5**), we conclude, this is a antigen specific effect. In contrast to the IAV response the antibacterial host profile was however largely characterized by proinflammatory marker genes (e.g. IL1A IL1B, TNF, IL6, IL17A) and interleukin signaling in the Reactome analysis (**Fig. 6D-F**). Lastly, we tested cell-to-cell communication in the human *ex vivo* model by blocking IL-1β signaling with a specific antibody. We compared the expression of IL-1β dependent genes after challenge with Spn in hPCLS. Without affecting the bacterial replication (**Fig. 6G**), the blocking of IL-1β signaling reduced the respective antimicrobial tissue response (**Fig. 6H**), confirming a robust cell-to-cell communication in this complex *ex vivo* model.

**Figure 6:**
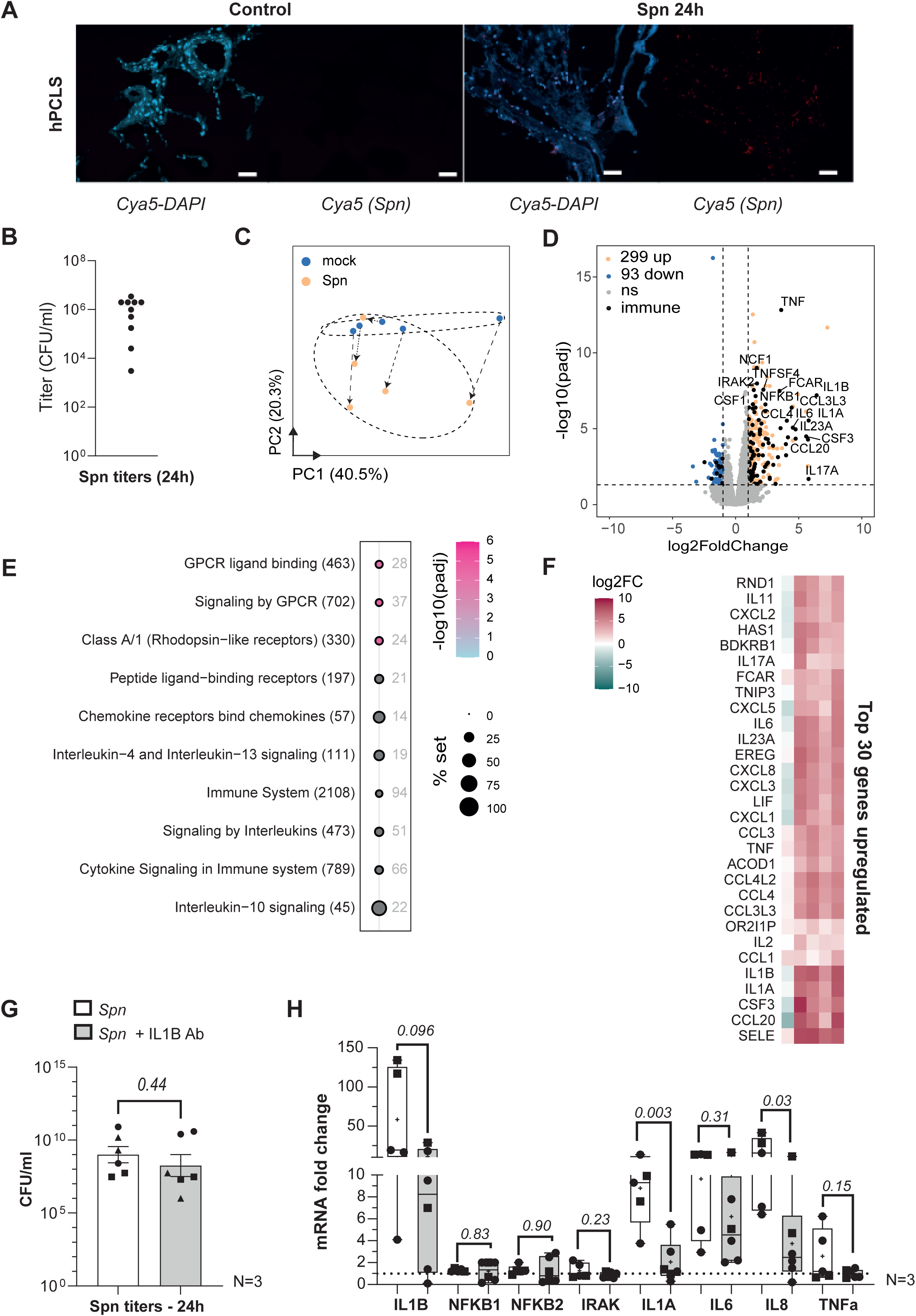
hPCLS can reproduce a classical antibacterial response towards a bacterial challenge. Human PCLS from five patients were infected with 10^3^ cfu of *Streptococcus pneumoniae* (Klein Chester ATCC 6303, serotype 3) for 24h. Bioinformatic analysis was done using Deseq2 with R programming. A) In situ RNA hybridization of dnaA (Cyanine 5) was used to visualize *Spn* at 24hpi in hPCLS sections. Magnification x100. Size bar is representing 100 µm. Mock are uninfected hPCLS. B) Bacterial titers from tissue homogenates of PCLS were measured at 24hpi (N=5, duplicates). C) Principal component analysis illustrates the global transcriptomics response of mock treated (blue) and Spn infected 24hpi (orange) hPCLS. Arrows indicate the corresponding infected sample for each patient mock sample. D) Volcano plot showing differentially expressed genes between Spn infected and mock treated samples. Log2 fold-change induction is indicated in the x-axis, p-adjusted in the y-axis. Genes have been represented in: orange (upregulated), blue (downregulated), grey (not significant). Genes associated to immune system pathways are represented in black. E) Pathway enrichment analysis was conducted using Reactome database (hypergeometric test). The top 10 pathways upregulated by *Spn* infection are shown. -log10 (p-adjusted) is given by a color scale. Coverage percentage of the pathway set is indicated by the size of the dot. Pathways are indicated with the total number of genes in brackets and the number of genes significantly regulated are indicated next to the dot. F) Heatmaps of the top 30 genes significantly upregulated (pink) and downregulated (blue) by Spn in hPCLS from 5 patients. Genes are ranked by the log2 Fold Change. G-H) hPCLS have been infected 24h with Spn (10^3^ cfu) and co-treated with an IL-1β blocking antibody (10µg/ml) (N=3, duplicates). G) Bacterial titers have been measured. H) RT-qPCR has been realized on a panel of genes associated to IL-1β signaling for Spn infection. Statistical significance was established with a one-way ANOVA (N=3, each symbol represents one independent experiment). White boxes are the Spn-infected hPCLS, grey boxes are co-treated with anti-IL-1β cocktail.

## Discussion

While primary respiratory tract epithelial cells have been shown superior to immortalized lung epithelial cells with regards to antiviral responses^16^, they are li Human lung-on-a-chip models, lung organoids or stratified airway epithelial cells (often used in air liquid interface) are mostly limited to the contribution of epithelial cells to lung infection biology. The PCLS model gains rapidly importance in biomedical studies targeting lung pathologies and pharmacological testing. mPCLS and hPCLS have been used to study the pathophysiology of inflammatory lung diseases^17,18^ and to test their response to disease relevant proinflammatory stimuli^19^. In respiratory infectious disease research animal PCLS and hPCLS gain importance, e.g., to assess the action of antimicrobials in a relevant organ context ^20,21^. Others used PCLS models before to assess the innate immune response to viral challenge^22–24^ or to colonization with different mixes of commensal bacteria (mPCLS)^25^. Two recent studies investigated viral replication and antiviral host responses in hPCLS by single cell RNAseq from a single human donor^26,27^. *Ex vivo* culture lung tissue from COPD patients with terminal lung emphysema was used to characterize early antiviral and antibacterial responses, 24h after IAV, *Pseudomonas aeruginosa* or *Mycobacterium tuberculosis* (BCG) challenge, by scRNAseq and bulk RNAseq^27^. Importantly, the authors noted a high donor dependent variation in host responses. They further stated that the pre-inflamed state of these tissues might not be fully representative of the antimicrobial response in healthy lung tissue. This could explain why only a limited number of IAV upregulated genes were found as compared to our study. We provide here comprehensive transcriptome data in human and murine ex vivo tissue compared to *in vivo* host responses and complement these findings with a detailed characterization of the cellular composition of the two PCLS models. While not surprising we show a clear contribution of the tissue resident memory cells to human host responses against IAV infection *ex vivo*. As for laboratory mice derived PCLS, the biological differences to hPCLS in responsiveness to microbial infection stem likely from different genetics, metabolism, microbiome and baseline immune status. The last point is of particular interest when studying infectious agents since many in vivo studies rely on immunologically naïve animals. Our data indicate that mPCLS from naïve mice rather poorly respond to IAV infection as compared to *in vivo* infections or hPCLS. In part the lack of infiltrating immune cells into the PCLS model explains these differences, but also the absence of tissue resident immune cells. Beyond this, especially infiltrating monocytes and neutrophils are contributing substantially to the cytokine response and eventually immune pathology observed during a severe viral pneumonia^28^. Hence proinflammatory signaling was not faithfully reproduced especially in the naïve murine *ex vivo* model. Exogenous addition of monocytes from blood or bone marrow might be a strategy to compensate for the absence of a connection to the supply with blood derived immune cells to overcome the limited antiviral response. PCLS from animals with multiple IAV encounters could serve as a better model to mimic the immune history observed in human patients, who on average undergo an IAV infection every 5-10 years^13^. Surprisingly, the innate antiviral host response did not differ in these animals. We found for example no increased expression of IFNγ, suggesting that tissue resident T-lymphocytes do not contribute to the increased protection. Considering the high number of B lymphocytes in mPCLS, we suspect an antibody mediated partial neutralization of viral infection. Globally, we observed specific antiviral and antibacterial responses in both PCLS systems, suggesting a clear distinction between the two types of pathogens. It must be noted that the timing (48h IAV *vs* 24h Spn) could in part explain these differences. Notably, a recent report indicated that PCLS undergo a spike of innate immune and inflammatory gene induction on the day of cutting which wanes within 24h^29^. It is unclear how this mechanic induction of the innate immune response affects a subsequent challenge with viral or bacterial pathogens. The lack of IFNγ induction by IAV infection in mPCLS was suggesting that the early antiviral response in *ex vivo* tissues naïve animals relies purely on a type I interferon responses. In contrast antibacterial/proinflammatory signaling after Spn challenge was readily detectable in mPCLS but less in hPCLS. This also suggests that mPCLS are not generally unresponsive to microbial exposure. In fact, an important contributor to the antibacterial host responses in mice is age. PCLS from aged mice display a stronger inflammatory reaction to bacterial PAMP than tissues from young animals^30^. In hPCLS from aged patients, we detected a distinct proinflammatory host response after Spn colonization, as compared to the antiviral response. Globally this response was less pronounced as the antiviral response despite robust bacterial replication. In fact, hPCLS from one human patient did not upregulate innate immune response genes after Spn challenge, but at the same time responded well to IAV, which suggest potentially an individual response based on previous exposure to pneumococci and potentially depending on patient specific subsets of tissue resident immune cells. Alternatively, tolerance mechanisms towards this common pathobiont in the upper respiratory tract of humans might play a role in dampening the antibacterial response. This could indicate that hPCLS could reflect the individual exposure history of a given patient and hence be used to study the basic cellular response in lung tissue based on immune history shaped cell composition. Such a tool would be useful to predict innate responses in lung tissue of different age groups, e.g., to novel influenza viruses, where preexisting immunity can play an important role to prevent mortality in critical target groups as reported for the 2009 H1N1 pandemic^31^. IFI27 for example was previously identified as marker for IFN responses in peripheral immune cells (monocytes and conventional DC) of influenza virus infected human patients^32^, suggesting that some of these immune cells could reside in hPCLS. The shortage of human biopsy/autopsy material collected early during an IAV infection makes it difficult to conclusively answer these questions. On a more applied note, we think that hPCLS could be used to test antivirals or anti-inflammatory drugs in a realistic environment with additional viral restriction provided by the tissue resident immune cells and with an architecture and accessibility reflecting that of the native tissue.

## Material and Methods

### Ethics

Human tissue was obtained from the thorax surgery of the university hospital Geneva (HUG) and its use was approved under the license 2022-01942 by the CCER. Written patient consent was provided. Animal experiments were approved by the cantonal authorities and institutional animal welfare board under license number: GE216).

### PCLS generation

mPCLS: C57BL/6J mice (female, 7–10 weeks of age) were purchased from Charles River Laboratories (France) and housed at least 7 days under SPF conditions with a strict 12h/12h light/dark cycle and food and water at libitum. Animals were euthanized by intraperitoneal injection of pentobarbital (150µg/g, Esconarkon®, Streuli Pharma, Uznach, Swtizerland) when experimental or humane endpoints were reached. To collect lung tissues, the abdominal cavity and the thoracic cage were opened surgically. The inferior vena cava was nicked, and lung lavage was performed by PBS 1X injection into the heart right vena. Next, a catheter (22G) was inserted in the trachea to perfuse the lungs with 2% low-melting-point agarose warmed to 37°C (Sigma-Aldrich, A9414, diluted in DMEM/Ham’s F12, Gibco). The mouse was cooled at 4°C for 10 min, then the lung was excised and incubated 30 min at 4°C to let the agarose solidified. The lobes were dissociated and murine PCLS (250µm thickness) were cut with a Krumdieck tissue slicer (Alabama R&D, USA, MD600).

hPCLS: The use of human tissue was approved by the cantonal ethics committee of Geneva (CCER, project number 2022-01942). Lung tissues were provided by the thorax surgery unit of the University Hospital Geneva (Hôpitaux Universitaires de Genève, HUG). Tumor-free tissue from lung cancer resection surgeries of 22 patients were collected for this work. The average age was 67.4 (±10.7) years, 12 were females and 10 males. Concerning smoking habits: 8 were active, 7 formers smokers, 3 non-smokers, and 4 were without information. Location of the resected tissue was in lower left lobe (3), lower right (2), upper left (6), upper right (8), middle right (3). Individual underlying disease state and medication are indicated in **Table S1**. The lung segment was washed with PBS 1X and perfused with 2% low-meting-point agarose thanks to catheters (22G). The solidification was allowed by 1h30 at 4°C on ice. Using a coring press (Alabama R&D, USA MD500), tissue cylinders of 8mm diameter were generated and transversally sliced into PCLS as described for mPCLS. PCLS were maintained in 500µL DMEM/Ham’s F12 (Gibco, supplemented with 100 U/mL penicillin and 100 µg/mL streptomycin (Sigma-Aldrich, P0781) without FCS) in 24-wells plate at 37°C, 5% CO2. The medium was changed 2 hours after cutting to remove the agarose, experiments were performed the day after the cutting to allow tissue recovering and agarose dissolution. Tissue integrity was confirmed with a brightfield microscope (Evos microscope system, Thermo Fisher).

### Bacteria

Spn (ATCC: (Klein) Chester 6303, serotype 3) was grown in Trypticase Soy Broth (Oxoid, UK, BO0369M) at 37°C with 5% CO_2_ in static culture up to 0.4 optical density (OD600 nm). PCLS infection was performed using 10^3^CFU per well in the PCLS culture medium at 37°C, 5% CO_2_. The inoculum was removed after 4h incubation, PCLS were washed two times with PBS 1X and fresh culture medium was added. The infection was stopped 24h post-inoculation. Supernatants and tissues were collected. Tissues were homogenized with a BeadBlaster homogenizer (Benchmark, D2400) using 1.4mm ceramic beads to determine bacterial titers (Omni International, 19627). Serial dilutions were spotted on Trypticase Soy agar plates with 5% sheep blood (BioMerieux, M1006) at 37°C with 5% CO2.

### Viruses

A/Netherlands/602/2009 (H1N1)^33^ was kindly provided by Dr. Florian Krammer (Icahn School of Medicine at Mount Sinai, New York, NY, USA). PCLS were infected with 10^5^ PFU in 200µl of PCLS medium per well. After 50 min of viral adsorption at 37°C, PCLS were washed with PBS 1X and incubated in fresh culture medium. Supernatants were collected at the indicated times post infection and viral titers were determined by plaque assay in MDCK cells (Madin-Darby canine kidney, ATCC).

### Plaque assays

MDCK cells were grown in DMEM (Dulbecco’s Modified Eagle Medium, Gibco) supplemented with 10% (vol/vol) heat-inactivated fetal bovine serum (Gibco) and 100 U/mL penicillin and 100 µg/mL streptomycin (Sigma-Aldrich, P0781). Cells (6-well plate format, 1,5 x 10^6^ cells/well) were infected with 10-fold serial dilutions in PBS 1X 0,2% BSA of the virus for 50 min at 37°C. Supernatants were replaced by fresh overlay medium supplemented with 1 mg/mL of N-tosyl-L-phenylalanine chloromethyl ketone (TPCK)-treated trypsin (Sigma) and 0,6% purified agar (Oxoid) and plates were incubated at 37°C 5% CO_2_. After 48 h, cells were fixed for 1 h at RT with 4% paraformaldehyde (PFA) and the overlays were removed. Cells were stained using a solution of 16% Methanol and crystal violet and plaques were counted visually.

### In vivo infection

For inoculation with IAV, mice were injected intraperitoneally with a mix of ketamine/xylazine (100 mg/kg and 5 mg/kg, respectively) in 200 µl of sterile PBS. Upon reaching deep anesthesia, mice were inoculated with 40 µl of PBS or virus (10^4^ PFU) via the intranasal route. Lungs were collected after 24h and 48 hpi for IAV.

### RNA extraction & RT-qPCR

Tissue was stored at −80°C in RNA protect Tissue reagent (Qiagen, 76104, Germany). PCLS were disrupted in the lysis buffer from the kit with 1.4mm ceramic beads (Omni International, 19627) using a BeadBlaster homogenizer (Benchmark, D2400). Then, RNA extraction was performed with the NucleoSpin RNA XS kit (Macherey-Nagel, 740902), following the manufacturer’s instructions. Elution was done in 30µl of water. RNA concentration was estimated by Nanodrop. 250 ng were used for reverse-transcription using the MMLV-RT (Invitrogen, 28025013). Gene expression was assessed by quantitative PCR using 2X KAPA SYBR FAST qPCR Master Mix-Universal (KAPA Biosystems, USA) and 10 µM of forward and reverse primers (**Table S2**). PCR was realized with CFX Connect Real-Time PCR detection system (Biorad) following this thermal cycling protocol: an initial denaturation step at 95 °C for 10 min, followed by 45 cycles of denaturation at 95 °C for 15 s, annealing/extension at 61.5 °C for 60 s, with a final melting curve step from 60 to 95 °C with 0.5 °C increment. The relative gene expression was calculated with the ΔΔCt method, using *hprt* (murine tissue) or 18s-rRNA (human tissue) genes for normalization.

For all lungs from mice or unprocessed lung biopsies from human, total RNA was extracted using TRIzol (Ambion, Life technologies) following manufacturer’s instructions.

### Lactate dehydrogenase (LDH) assay

To determine global cell suffering, LDH release was quantified daily into the PCLS supernatant using the colorimetric test CYQUANT ^™^ LDH cytotoxicity assay kit (Invitrogen, C20300), according to the manufacturer’s recommendations.

### RNAscope

PCLS from mice and human were collected under sterile conditions, fixed in PBS 1X - paraformaldehyde 4% solution overnight at 4°C and embedded in paraffin and sliced using a standard microtome (5 µM thickness). *In situ* detection of *Streptococcus pneumoniae* (RNAscope® Probe - B- S. pneumoniae -SPN23F12870-C2, accession number: FM211187.1, nucleotides: 2-753) and IAV (RNAscope® Probe V - Influenza A - H1N1 – segment 5 – NP - O8 -C1, accession number: CY176945.1) was performed using the RNAscope 2.5 HD Assay - RED (Advanced Cell Diagnostics) according to the manufacturer’s protocol. Briefly, tissue slices were deparaffinized by 1h incubation at 60°C followed by four rounds in 100% xylol and four rounds in 100 % ethanol solutions. An incubation with hydrogen peroxide was performed for 10 min and target retrieval was achieved by incubating the slides with RNAscope 1x Target Retrieval Reagent for 45 min at 95°C. Tissue was permeabilized using RNAscope protease plus for 30 min (Advanced Cell Diagnostics). Probe hybridization was performed with a specific RNA probe for IAV or Spn for 2 h at 40°C. Signal amplification was achieved by incubating the tissues at 40°C with AMP solutions provided by the kit. Cyanine 3 and cyanine 5 were used as fluorophores respectively for IAV and Spn. Slides were counterstained with Prolong Diamond Antifade Mountant with DAPI (Invitrogen, P36962). Slides were visualized under Nikon A1r spectral microscope under a x10/20 objective and processed using Zen Software.

### Transcriptomic analysis

RNA was extracted as described before at the replication peak of each pathogen (48h for IAV, 24h for Spn). RNAseq (30 million reads per sample (triplicates for each murine condition, five patients samplings for hPCLS) was performed at the iGE3 Genomics Platform at the University. Analysis of these data was realized with the help of the Bioinformatics Support Platform (UNIGE). Libraries were prepared with the Illumina TruSeqHT Stranded mRNA protocol for PCLS and mouse in vivo lungs and with RNA exome-seq for human in vivo autopsy samples and corresponding control. The sequencing was done on Illumina NovaSeq 6000 sequencer. Sequenced reads were mapped with HISAT2 aligner either to the mouse genome GRCm39 (source: ensembl 110) or to the human genome GRCh38.p14 (source: ensembl 110). Read-count quantification was extracted with the method summarizeOverlaps() from R package GenomicAlignments. The code and softwares used in the analysis were packaged into a docker container made publicly available at https://github.com/BioinfoSupport/ngs (container), https://github.com/ BioinfoSupport/rnaseq (mapping and quantification pipeline), and https://github.com/BioinfoSupport/genomes (for HISAT2 indexed genomes). Normalization, differential gene expression analysis, and data visualization were conducted with the R programming language. Package DESeq2 was used to asset the statistical significance of differential gene expressions. More specifically, we conducted 4 independent DESeq2 analyses to take into consideration samples pairing: A) one with 6 unpaired samples of the mouse in vivo conditions (mIV_mock, mIV_IAV); B) another with the 6 paired-samples of the mouse PCLS conditions (mPCLS_mock, mPCLS_IAV); C) a last with 15 paired samples of the human PCLS conditions (hPCLS_mock, hPCLS_Spn, hPCLS_IAV); D) one with 6 unpaired samples of the human in vivo conditions (hIV_mock, hIV_Influenza). The details of the samples used in each set is given in **Table S3** “RNA-seq Samples description”. Pathway enrichment analysis was done under the R programming language with hypergeometric tests against Reactome.org database. Transcription factor enrichment analysis was realized using EnrichR^8–10^.

### Flow cytometry

One day after slicing, 14 PCLS per condition were washed with cold PBS 1X and pooled together. Cell dissociation was performed for 45 minutes at 37°C (aggitation) with PBS +5% FBS with 0.5 mg/mL Collagenase D and 0.1mg/mL DNaseI. Cells were centrifugated 5 min at 300g. Red blood cell lysis was performed with ACK buffer for 4 minutes at RT. The reaction was stopped with RPMI 10% FBS and cells were spined and washed in PBS.

Cells were stained 15 min at RT with Fixable Viability Dye (Invitrogen, 1:1000 in PBS 1X) and FC blocking was performed with 0.25µg of TruStain FcX PLUS (Biolegend) in 100µL of MACS before staining. Cells were then stained 15 min on ice with surface markers antibodies resuspended in MACS buffer (PBS-0.5% BSA-2mM EDTA) according to the panel. Antibodies references are indicated in **Table S4**. Cells were fixed with 200 μL of IC Fixation Buffer (Thermofisher) for 30 min at RT. Samples were centrifugated at 450g at RT for 5 min. Cells were resuspended in an appropriate volume of MACS buffer and the samples were run on LSRII Fortessa instrument (BD Biosciences).

### IL-1β blocking experiment in hPCLS

hPCLS were infected with 10^3^ CFU of Spn, after 4h of attachment, PCLS were washed 2 times with PBS 1X and culture medium containing 10µg/ml antibody anti-ILβ (R&D Systems, MAB601, clone 2805) was added. Tissues and supernatants were collected 24hpi. Stastistical analysis was

### Statistical analysis

Statistical analysis was performed using GraphPad Prism 9 or R and statistical tests applied as well as the number of biological and technical repeats are indicated in each respective figure legend.

## Acknowledgments

Our work was supported by a generous donor, advised by CARIGEST SA and the Fondation privée des HUG. We are extremely grateful for the generous donation of tissue samples by HUG patients in support of our research and for the professional collaboration with the surgical team of the HUG providing this. We would further like to thank the animal care takers at the CMU for their daily support as well as the core facilities for iGE3 Genomics, Bioimaging, Histology and Cytometry. We are grateful for the insightful advise by Prof. Dorothee Viemann and Prof Benjamin G. Hale.

## Supplementary figures

**Figure S1: Viability of mPCLS and hPCLS in culture.**

Cell death was measured in mPCLS (in green) and hPCLS (in gray) by quantification of LDH release upon 4 days post-slicing. Positive control of cell death (100%) was realized with triton 10X treatment (N=3, duplicates).

**Figure S2: Flow cytometry gating strategy to describe immune cells in murine PCLS**

Examplary gating strategy is shown for A) Lymphoid cell panel B) Myeloid cell panel

**Figure S3: Signaling activation in mouse in vivo and ex vivo 48h after IAV infection**

Mice and mPCLS were respectively infected with 10^4^/10^5^ PFU of IAV (A/Netherlands/602/2009/H1N1) for 48h. Tissue was processed to bulk RNA sequencing and host response was analyzed using ‘DeSeq2’ package on R. Transcription factor enrichment analysis was conducted using EnrichR platform for IAV host responses in mouse in vivo (A) and ex vivo (B). Only transcription factors significantly enriched are indicated. Odds ratios are indicated in x-axis and p-value in y-axis. Hemostasis pathway (C), Extracellular matrix organization (D) and Programmed cell death (E) heatmaps in mouse system 48h after IAV infection

**Figure S4: Flow cytometry gating strategy to describe resident memory lymphoid cells in murine PCLS**

Examplary gating strategy is shown for a lymphoid cell panel

**Figure S5: Flow cytometry gating strategy to describe immune cells in human PCLS**

Examplary gating strategy is shown for A) Lymphoid cell panel B) Myeloid cell panel.

