## Supplementary figures and images for "*Ex vivo* modelling of lung tissue resident antimicrobial responses"

### fig s1

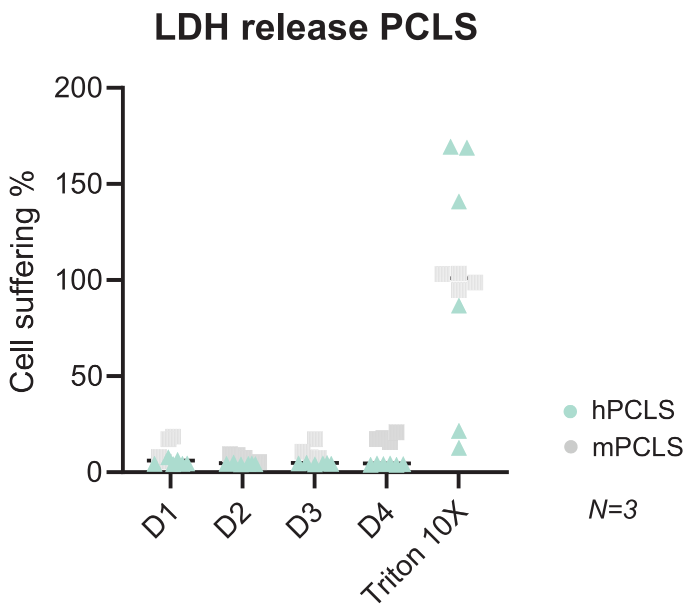

### fig s2

**A**

##### Lymphoid gating strategy: mouse

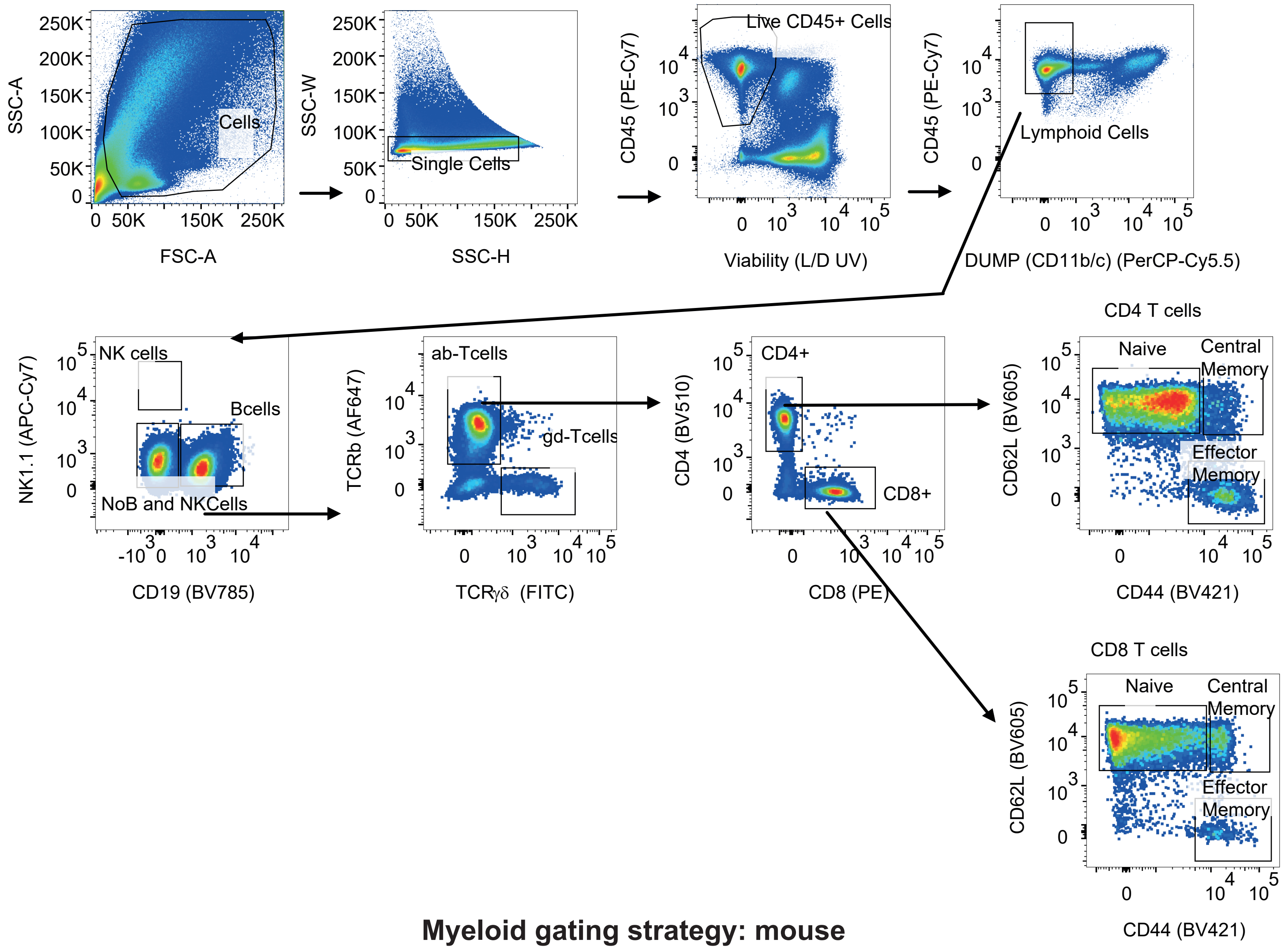


**B**

##### Myeloid gating strategy: mouse

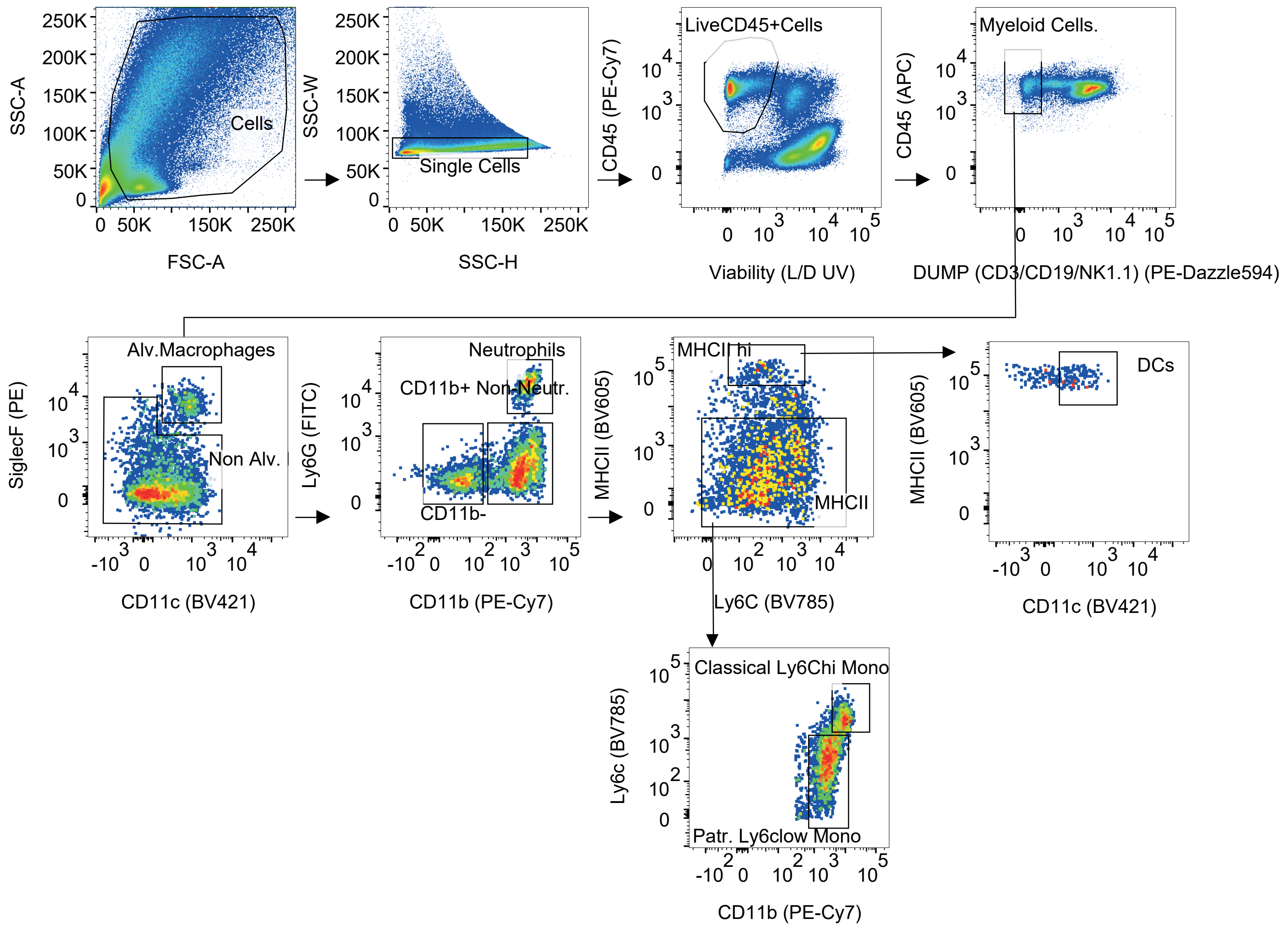

### fig s3

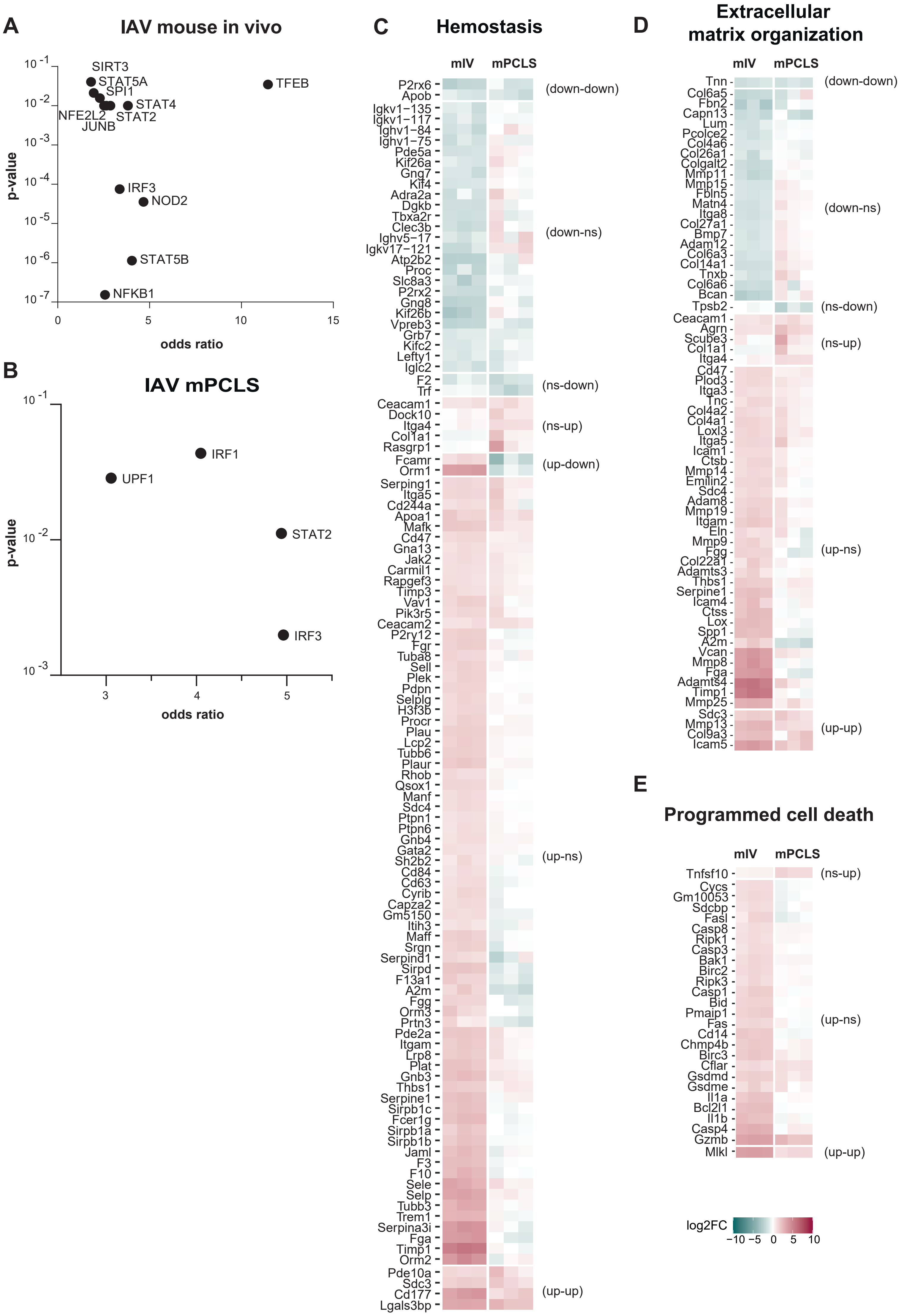

### fig s4

#### Gating strategy - priming experiment - murine tissue

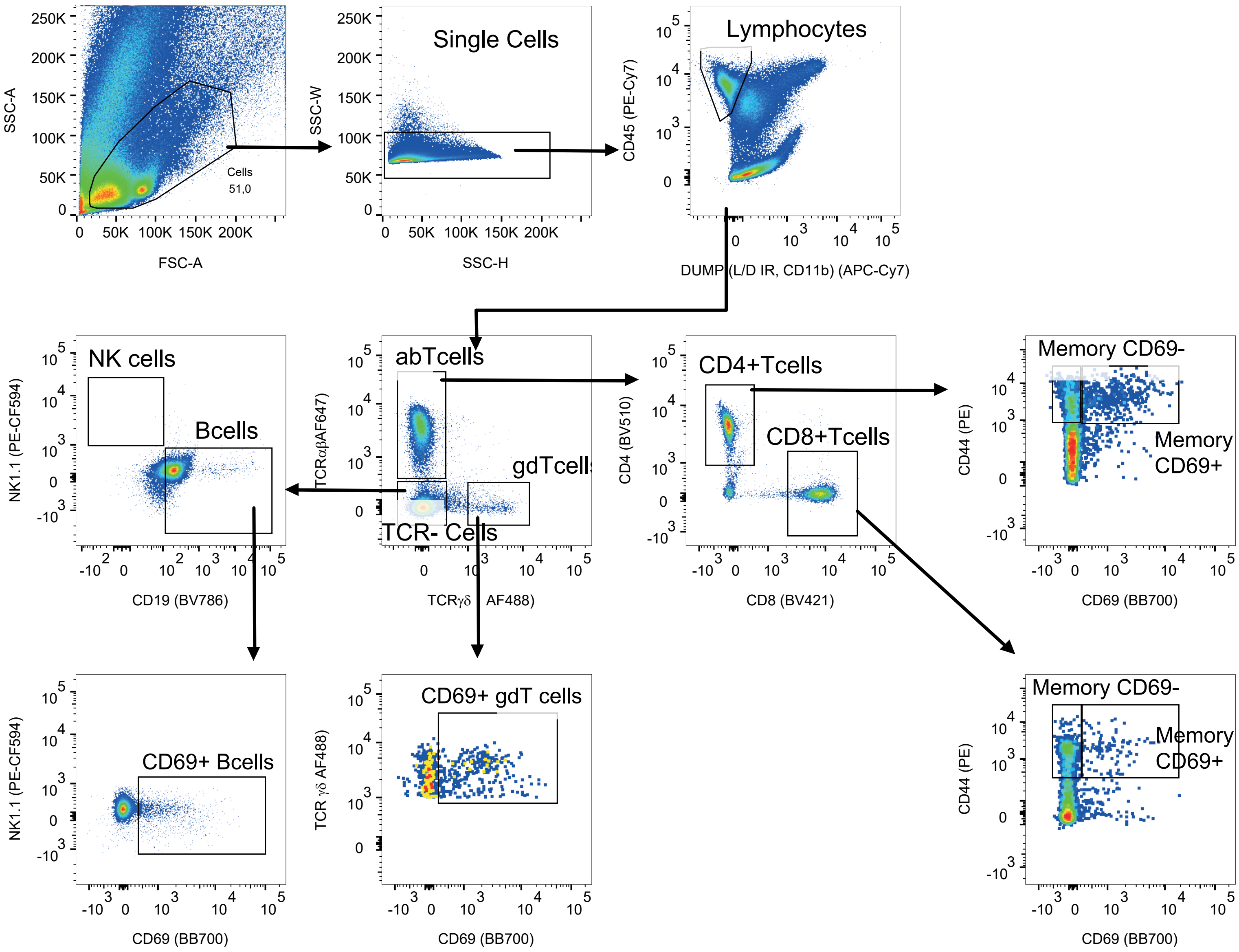

### fig s5

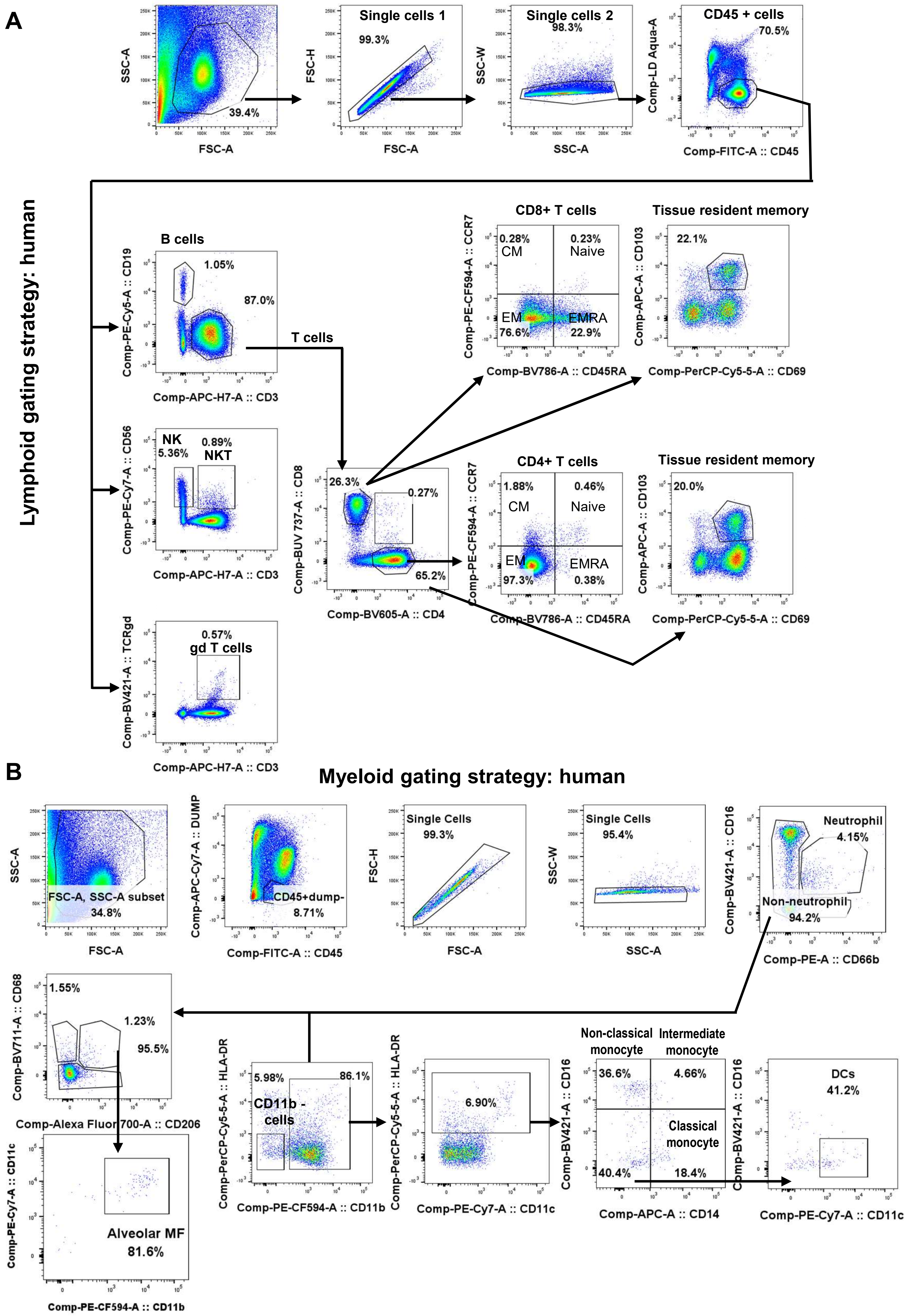
